# Influence of effective polarization on ion and water interactions within a biomimetic nanopore

**DOI:** 10.1101/2021.12.10.471283

**Authors:** Linda X. Phan, Charlotte I. Lynch, Jason Crain, Mark S.P. Sansom, Stephen J. Tucker

## Abstract

Interactions between ions and water at hydrophobic interfaces within ion channels and nanopores are suggested to play a key role in the movement of ions across biological membranes. Previous molecular dynamics (MD) simulations have shown that the affinity of polarizable anions to aqueous/hydrophobic interfaces can be markedly influenced by including polarization effects through an electronic continuum correction (ECC). Here, we designed a model biomimetic nanopore to imitate the polar pore openings and hydrophobic gating regions found in pentameric ligand-gated ion channels. MD simulations were then performed using both a non-polarizable force field and the ECC method to investigate the behavior of water, Na^+^ and Cl^−^ ions confined within the hydrophobic region of the nanopore. Number density distributions revealed preferential Cl^−^ adsorption to the hydrophobic pore walls, with this interfacial layer largely devoid of Na^+^. Free energy profiles for Na^+^ and Cl^−^ permeating the pore also display an energy barrier reduction associated with the localization of Cl^−^ to this hydrophobic interface, and the hydration number profiles reflect a corresponding reduction in the first hydration shell of Cl^−^. Crucially, these ion effects were only observed through inclusion of effective polarization which therefore suggests that polarizability may be essential for an accurate description for the behavior of ions and water within hydrophobic nanoscale pores, especially those that conduct Cl^−^.

## Introduction

The interactions of ions and water with membrane-embedded nanopores is of significant biological and technological importance. Nanopores in technology span a wide range of applications including water desalination, DNA sequencing and biosensing, all of which exploit their ability to conduct and often differentiate between charged ions (1, 2). In biology, ion channel pores typically have an internal radius of ~ 0.5 nm and a length of ~ 3 nm. They are responsible for enabling and regulating the movement of ions across lipid bilayers and so can be considered as nanoscale devices (3).

Physiological processes rely on their correct functioning, and many diseases (channelopathies) result from their malfunction (4). The importance of understanding both biological ion channel and synthetic nanopore function has therefore led to a sustained interest in the molecular behaviour of ions and water in such nanoconfined environments (5). Yet despite many previous studies, a universal consensus on these interactions has not yet been reached, and is partially due to contradictory trends in the many energetic contributions to anionic and water interactions with hydrophobic interfaces (6, 7).

Ion and water permeation through subnanometer pores is influenced not only by pore radius, but by the local hydrophobicity of the pore lining. Permeation may readily occur through polar regions with dimensions just larger than the radius of the permeating species (8). However, for hydrophobic regions of comparable dimensions, nanopores may spontaneously dewet, leading to an associated energetic barrier to permeation without steric occlusion (8, 9). This concept is referred to as hydrophobic gating (8, 10, 11) and has been demonstrated in both biological (10, 12–15) and synthetic nanopores (16, 17). A number of recent structural studies have also indicated that hydrophobic surfaces within channels and nanopores may provide favorable interaction sites for anions such as Cl^-^ that can influence the functional properties of these pores (18, 19). Therefore, an accurate description of the interactions of ions and water with hydrophobic interfaces is essential for the understanding of ion permeation in the confined environments found in such pores.

Extensive studies have been carried out on electrolyte solutions at hydrophobic interfaces such as the aqueous/air interface (6) where molecular dynamics (MD) simulations employing explicitly polarizable force fields have revealed a propensity for halide ions to associate at the interface following the Hofmeister series, i.e. F^-^< Cl^-^< Br^-^< I^-^. This order is directly correlated to the polarizability of the anion so that smaller ‘hard’ F^-^ ions are excluded from the aqueous/air interface, whereas the larger polarizable halides demonstrate an increasing affinity towards the interface (6, 20–22). This phenomenon can be explained in terms of the polarizable anion dielectric continuum theory (PA-DCT) which considers the hydration and polarizability of the anions (23, 24). The PA-DCT suggests that there is an attractive interaction between the induced dipole in larger weakly hydrated anions with surrounding water molecules at the interface that compensates for the loss of water-ion hydrogen bonds (6, 25). Therefore, large polarizable anions are capable of partially losing some of their hydration shell whilst remaining stable at the aqueous/air interface. Similar effects have also been observed at more complex hydrophobic interfaces such as the aqueous/oil interface (26, 27). Notably, the anion of most biological significance, Cl^-^, falls in the middle of this series and so its behavior in simulations is likely to be sensitive to the treatment of such interfacial interactions.

Direct experimental observations of ion behavior in interfacial regions remains a challenge. However, interfacial ion properties can be inferred from spectroscopic techniques able to sample surface regions of electrolyte solutions, which can elucidate the hydrogen bonding environment and probe surface ion concentrations (6, 28, 29). These experimental results are largely in agreement with simulation, but nevertheless the quantitative extent of anion adsorption to the aqueous/air interface continues to be a matter of discussion (30).

MD simulations are therefore a useful tool to provide a molecular interpretation that complements such experimental measurements. Many MD studies of nanopores and interfaces employ classical, non-polarizable (NP) force fields which do not fully capture the electronic response to the local environment (31). Neglecting polarizability therefore has consequences for accurately modelling the properties of polarizable anions where many key effects of these ions arise from their electronic responses. For example, the use of NP force fields can lead to inaccuracies in describing short-range ion-water and ion-ion interactions as well as overestimating ion clustering (32, 33).

It has been suggested by Leontyev & Struchebrukhov (34), and expanded upon by Jungwirth *et al*. (26, 35–37), that the lack of polarization in NP force fields can be compensated for by implicitly accounting for electronic polarization effects through an electronic continuum correction (ECC). In this approach the integer charges on monatomic ions in aqueous electrolytes are rescaled by a factor of 1/*ε_el_*^½^ (26, 34, 38). Here, *ε_el_* represents the electronic component of the dielectric constant which can be taken as the high-frequency dielectric constant (*ε_el_* = 1.78 for water and *ε_el_* ~ 2 for proteins).

Thus, ECC may be considered as an approach to maintain consistency in the parameterization of ions, alongside that of water and biomolecules (e.g. proteins, lipids etc.) which are already routinely parameterized with partial charges to account implicitly for electronic polarization (5, 32). Aqueous/hydrophobic interfaces are especially suited to the use of ECC because they have an approximately uniform high-frequency dielectric constant across the system i.e., the values of *ε_el_* for each media are comparable (26, 37). Vazdar *et al*. demonstrated that when ECC is applied to an aqueous/oil interface, there is significant improvement in bulk aqueous salt solution properties relative to NP force field simulations, yielding simulation results in agreement with experimental findings and also comparable to explicitly polarizable force fields (26). These improvements to the ion force field are critical for studies concerning the dynamics of weakly polarizable anions, especially Cl^-^.

The precise mechanism that underlies Cl^-^ selectivity in nanopores and channels is also poorly understood, and Cl^-^ channels are often less intensively studied than their cation conductive counterparts. Unlike cation channels that often have high affinity and selectivity, Cl^-^ channels are often permeable to other anions (39). Furthermore, it is often unclear how interactions between Cl^-^ and hydrophobic contacts can influence Cl^-^ selectivity and consequently determine their functional properties.

With the emergence of many new structures for anion selective channels (40–43), it is therefore of particular importance to investigate the relationship between hydrophobic contacts and the dynamic behavior of ions and water within their pores. An improved understanding of such interactions will also facilitate design of biomimetic nanopores (44). Certain aspects of such pores can be effectively mimicked by simple non-biological structures, for example graphene nanopores and metal-organic structures (2). Carbon nanotubes (CNTs) are also particularly attractive as structural templates because they can imitate many fundamental aspects of such pores including high transport efficiency, tunable pore diameters, functionalization, and well-defined hydrophobic interiors (45–50). The relevant properties of CNT nanopores have been extensively studied, both experimentally and with simulations (49, 51–53).

Here, we designed a simple biomimetic nanopore to explore the dynamics of ion and water interactions under hydrophobic confinement. We then performed MD simulations of the model nanopore with a NP force field and a NP force field with ECC-rescaled ionic charges to investigate the localization of ions and water relative to the internal hydrophobic nanopore interface. Potential of mean force (PMF) calculations also allowed us to examine the free energy landscapes of ions along the long axis of the pore. Finally, we have explored the hydration structure around these ions at various locations within the central hydrophobic section of the pore. We are thus able to compare the behavior of ions and water inside the pore when modelled by a classical NP force field and for ECC-rescaled ionic charges. Our results demonstrate that modelling polarizability quantitatively alters our model of nanopore/ion interactions and will be important for our understanding of ion permeation in general, especially in Cl^-^ channels.

## Methods

### Nanopore Models

Pristine armchair (14,14) CNTs were generated and capped with hydrogen atoms to form the hydrophobic pore using the molecular builder, Avogadro (54), and VMD (55). The length of the CNT was ~ 4.7 nm (and therefore capable of spanning the thickness of the membrane) and the internal diameter was ~ 1.4 nm. A smaller armchair (10,10) CNT was built as a template for insertion and restraining of water molecules in selected positions to create the polar regions of the pore. A harmonic restraining potential was applied between the oxygen of the water molecules and the carbon atoms of the CNT pore wall interiors with a force constant of 1200 kJ mol^-1^ nm^-2^ and a maximum distance of 0.143 nm. For simulations investigating the effects of pore radius, (14,14), (16,16) and (18,18) armchair CNTs were used with internal radii of 0.70 nm, 0.83 nm and 0.95 nm respectively. (10,10), (12, 12) and (14,14) armchair CNTs were used as templates for water molecule insertion to create polar regions for the (14,14), (16,16) and (18,18) CNTs respectively. Pore radius profiles of the resultant model nanopores were calculated using the Channel Annotation Package (CHAP) (56).

### Molecular Dynamics Simulations

We performed 50 ns atomistic MD simulations of the model nanopore embedded in a POPC (1-palmitoyl-2-oleoyl-*sn*-glycero-3-phosphocholine) bilayer. The nanopore was inserted into the POPC bilayer by the InflateGRO method whereby the nanopore was placed in the membrane and equilibrated (57). The system was then solvated with a 0.50 M NaCl solution. All systems were first equilibrated for 10 ns and this period was not included in analysis. The simulations were carried out using GROMACS 2020 (www.gromacs.org) (58) with the OPLS all-atom force field with united-atom lipids (59) and the SPC/E water model (60). The integration timestep was 2 fs. All bonds were constrained using the LINCS algorithm (61). A Verlet cutoff scheme was applied and long-range electrostatics were treated by the particle mesh Ewald (PME) method (62) with a short-range cutoff of 1 nm and a Fourier spacing of 0.16 nm. Three-dimensional periodic boundary conditions were applied. Simulations were performed in the isothermal-isobaric ensemble. The temperature was maintained at 300 K with a coupling constant of *τ_t_* = 0.5 ps with a Nose-Hoover thermostat. Pressure was maintained semi-isotropically using the Parrinello-Rahman barostat at 1 bar with a coupling constant of *τ_t_* = 2.0 and compressibility of 4.5 × 10^-5^ bar^-1^. For simulations exploring the effects of ion concentration, NaCl concentrations from 0.25 M to 1.0 M were considered in 0.25 M increments. Electronic polarization effects were introduced to the system in a mean-field approach by applying the ECC method (26, 35–37). This was realized by rescaling all ionic charges by 1/*ε_el_*^½^ where *ε_el_* = 1.78 is the high frequency dielectric constant for water, thereby equating to a scaling factor of 0.75. In all simulations, the model nanopore was modelled using an additive force field. Three independent repeats were carried out for each individual parameter combination. Data were analyzed using GROMACS and locally written code using MDAnalysis (63–65).

### Umbrella Sampling

Umbrella sampling was performed to obtain one-dimensional potential of mean force profiles for Na^+^ and Cl^-^ ions moving through the model nanopore using both the non-polarizable force field and ECC method. Simulation details were similar to those detailed above. However, to prevent the nanopore from tilting in the bilayer, the carbon atoms of the CNT were placed under a harmonic restraint with a force constant of 1000 kJ mol^-1^ nm^-2^. Equilibration simulations were performed for 15 ns and the starting configurations for the umbrella windows were produced from the final state of these simulations.

The reaction coordinate was defined as the z-axis which corresponds approximately to the pore axis and direction normal to the lipid membrane. A target ion was relocated to subsequent positions along the z-axis followed by 10 steps of energy minimization to remove any steric clashes between the target ion and surrounding water molecules. A harmonic biasing potential was applied to the z-coordinate of the target ion with a force constant of 1000 kJ mol^-1^ nm^-2^. Umbrella windows covered the entire length of the model nanopore and up to 1 nm outside the pore. This setup corresponds to 69 windows along the z-axis with a distance of 0.1 nm between successive windows. Each umbrella window was simulated for 10 ns. PMF profiles were obtained through unbiasing with the Weighted Histogram Analysis Method (WHAM) using the Grossfeld lab implementation in version 2.0.9 (http://membrane.urmc.rochester.edu/wordpress/?page_id=126”). The first 5 ns of each simulation were discarded as equilibration, meaning that the final PMF profile was calculated from the final 5 ns of the simulation time. Each resulting PMF was zeroed with respect to the environment outside the nanopore. Convergence was assessed by comparing the cumulative free energy profiles computed from 1 ns fractions of simulation time (Fig. S8).

## Results & Discussion

### Designing a biomimetic nanopore

In designing a model nanopore, we sought to explore the interactions of Cl^-^ with hydrophobic interfaces in a simplified system representative of a biological ion channel. Based on concepts from our previous studies of pentameric ligand-gated ion channels (pLGICs) (12, 18), we set out to construct a model nanopore that could mimic the general charge distributions of different regions inside such channel pores. pLGICs are a family of ion channels that mediate fast neurotransmission (66). Within their pores, a highly conserved hydrophobic pore-lining region is associated with channel gating (12). There are also narrow regions near the entrances of the pore formed by polar and charged residues (10, 67, 68). We therefore mimicked these two regions in a model nanopore built from a CNT.

The polar regions at either mouth of the nanopore were constructed by applying a harmonic restraining potential between a set of water molecules and the interior pore walls near the openings whilst the central internal hydrophobic cavity was left exposed to resemble the hydrophobic gating region. The resultant model nanopore therefore has three defined regions of alternating polarity and hydrophobicity along its pore axis (**Fig. 1A,B**). This pore was then embedded into a phospholipid bilayer to span the thickness of the membrane (~ 4.7 nm) and form a stable membrane-embedded nanopore (**Fig 1C,D**).

**Figure 1:**
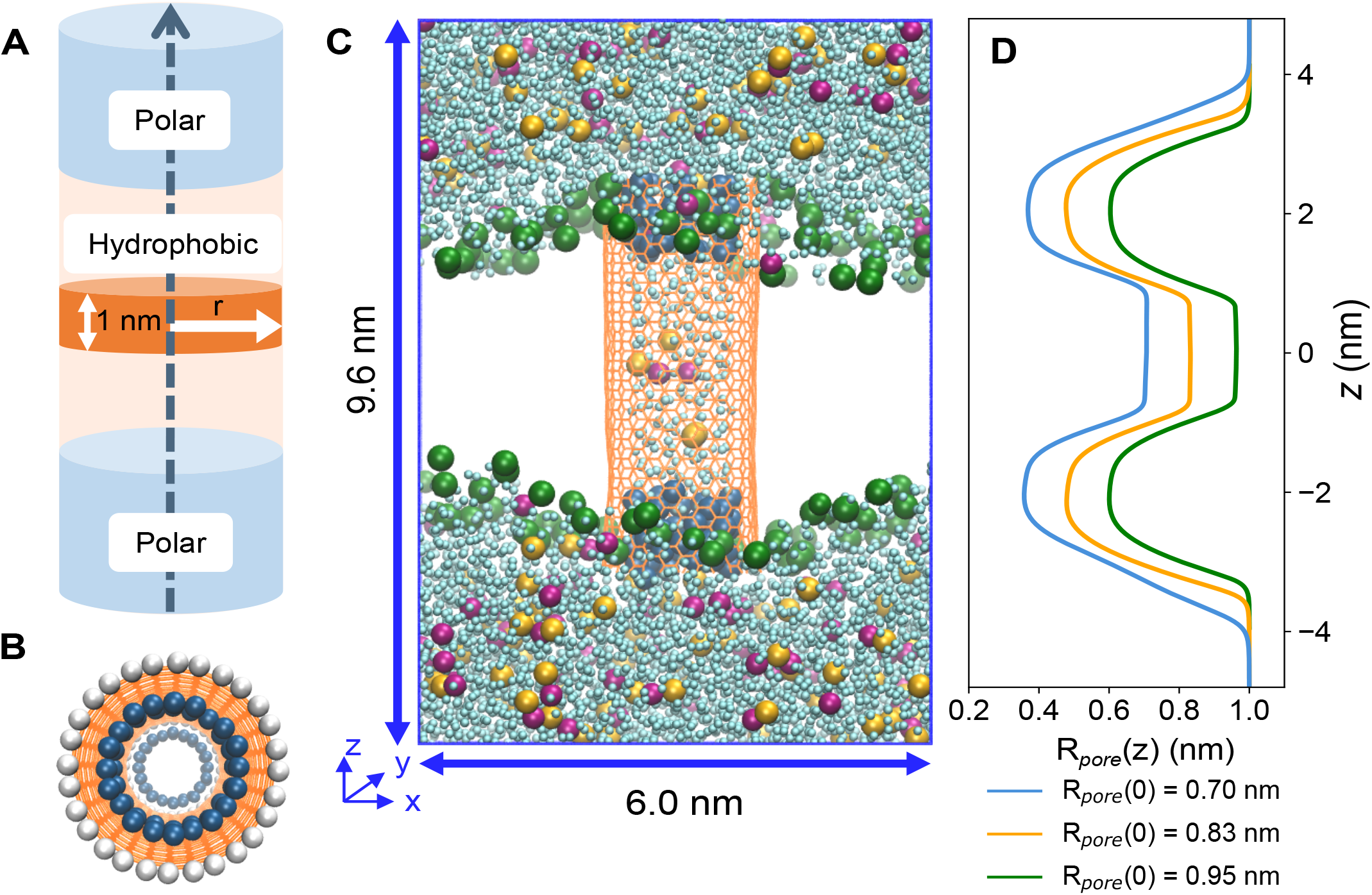
**A** Schematic of a biomimetic nanopore. The model nanopore has a radius r, such that 0 < r < R*_pore_*. **B** Top-down view of the pore, showing water molecules (blue) restrained to the CNT pore walls (orange) to create the polar regions at each mouth of the pore. The nanopore ends are capped with hydrogen atoms (white). **C** Snapshot of the simulation setup. The biomimetic nanopore is embedded in a POPC bilayer (lipid headgroup phosphates in green) and solvated with water (light blue), Cl^-^ (yellow) and Na^+^ (pink). **D** Pore radius profile showing the maximum value of R*_pore_* as a function of axial position *z* (approximately aligned with the simulation snapshot in **C)**.

### Influence of effective polarization

We performed two sets of simulations of model nanopores in the presence of NaCl solution: non-polarizable (NP) using the additive OPLS-AA force field (59) and ECC using the same force field with ECC-rescaled ionic charges. In both cases the SPC/E water model was employed (60). We then derived the number density profiles of the ions and water along and radially to the pore axis. The influence of the internal pore radius and NaCl concentration on the ion and water densities inside the hydrophobic central region of the pore was studied by sampling from a 1 nm thick slice along the z-axis i.e. from z = −0.5 to +0.5 nm for the final 10 ns of each 50 ns simulation.

Radial density profiles of ions when employing ECC-rescaled charges exhibited increased propensity Cl^-^ association with the hydrophobic pore lining, as indicated by a density peak displaced towards the water/CNT interface (**Fig. 2A**). By comparison, the Na^+^ ions are largely excluded from the immediate vicinity of this hydrophobic surface and are localized closer to the pore axis where they remain more fully solvated (**Fig. 2B**). Significantly, these ion distributions (which match those seen in simulations of ions in water nanodroplets (22, 69)) were only observed when implicitly including polarization through the ECC method. This is in marked contrast to simulations with the NP force field in which both anions and cations were equidistant from the hydrophobic pore wall, preferring to reside close to the pore axis where they are more fully solvated (**Fig. 2D & 2E**). These results reveal similar surface effects to those observed at the aqueous/decane interfacial system (**Fig. S1** in the Supporting Material) and are in good agreement with earlier studies of aqueous/air interfaces (21, 26, 35) as well as first principles MD and polarizable simulations of NaCl inside CNTs (70, 71).

**Figure 2:**
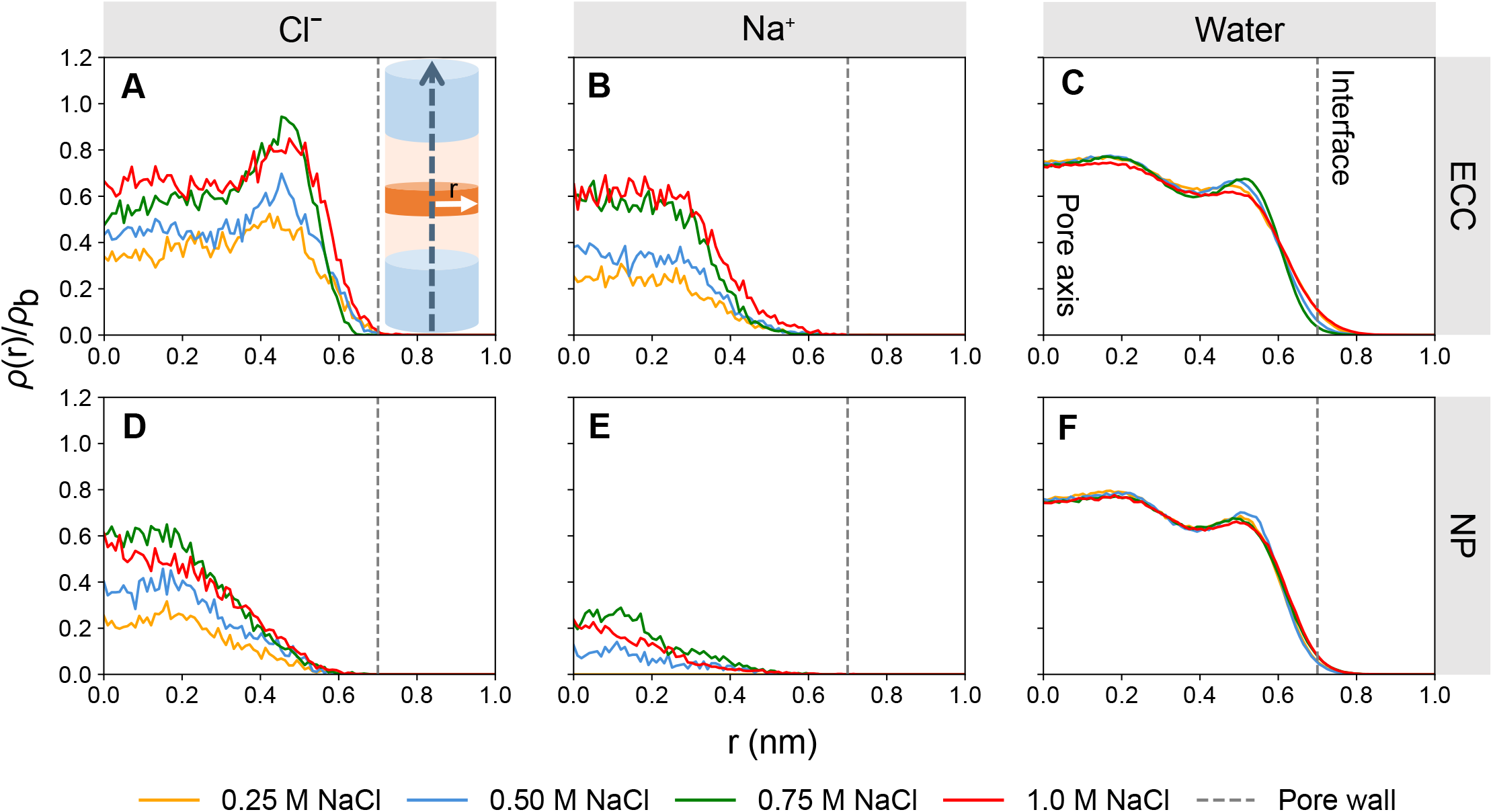
Symmetrized number density profiles of Cl^-^ (**A** & **D**), Na^+^ (**B** & **E**) and water (**C** & **F**), with ECC-rescaled ionic charges and a non-polarizable (NP) force field for all other atoms, at various salt concentrations. ρ(r)/ρ_b_ represents the symmetrized number density, ρ(r), sampled from the hydrophobic region of the nanopore (orange section of schematic (Fig. 1A)) and normalized by bulk density, ρ_b_. The variable r is the radius of the nanopore which extends from 0 (pore axis) to R*_pore_* (the interface where the salt solution meets the wall of the nanopore). The grey vertical dashed line represents R*_pore_*.

There is some degree of structuring of the ions and water inside these pores. Notably, the water density distributions (**Fig. 2C & 2F**) form a layered structure inside the nanopore with the outmost layer forming an ordered concentric ring and inner layers demonstrating more bulk-like profiles. In simulations with ECC, it is this outermost layer that is shared with Cl^-^ ions. Comparable structured behavior of water in concentric shells inside pristine CNT porins has previously been reported (16, 72, 73).

Given the pore is open, and ions and water are freely allowed to permeate, the area under the normalized number density curve indicates the total number of particles in the sampling region. At any given salt concentration, there are significantly fewer sodium ions present in the hydrophobic core of the nanopore relative to the number of Cl^-^ (**Fig. 2A,B,D,E**). This observation is more pronounced when applying the NP force field, with virtually no Na^+^ present in the hydrophobic core at lower concentrations, suggesting there is competition between Cl^-^ and Na^+^ to occupy regions close to the pore axis where water density is more bulk-like. With the improved electronic description using ECC, an unequal permeability ratio between ions persists, whilst interfacial properties are also captured. It is thought that ion pairs of unusually long lifetimes can form within these dimensions. However, our understanding of the cotransport of different ion species under nanometer confinement is incomplete (74).

### Cl^-^ accumulation at the hydrophobic pore wall

We next examined the behavior of ions and water inside model nanopores with different radii using number density calculations to explore the influence of different degrees of confinement (**Fig. 3**). To investigate this, model nanopores with radii (*R_pore_*) for the hydrophobic region of approximately 0.70 nm, 0.83 nm and 0.95 nm, were built. These pore dimensions are comparable to the hydrophobic gate of a simplified open-state β-barrel structure (16). All systems were simulated with 0.50 M NaCl solution. Analysis protocols were the same as those detailed for exploring ion and water number density as a function of salt concentration, however these data were now normalized to the surface area of the pore to focus on interfacial effects.

**Figure 3:**
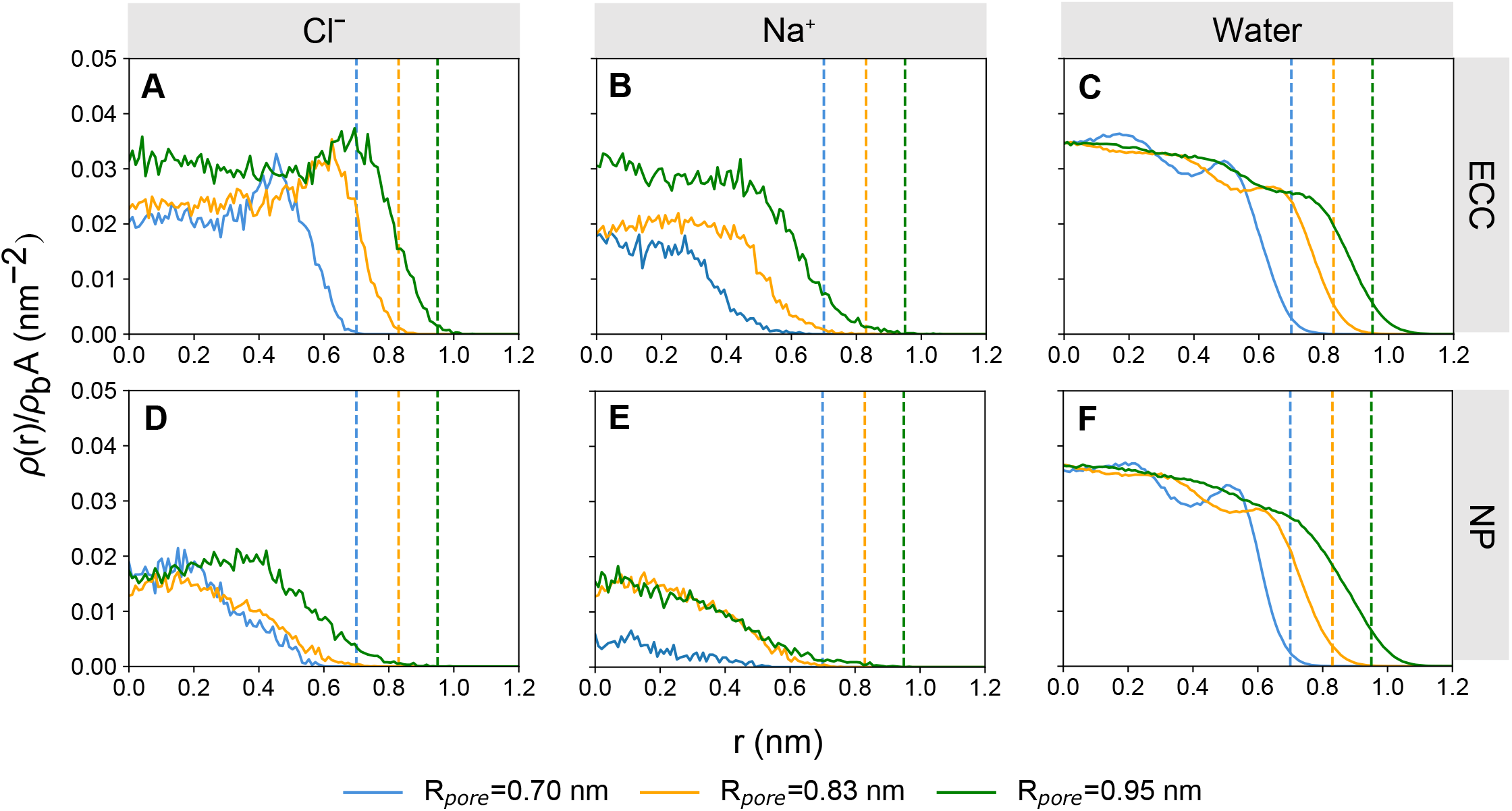
Symmetrized number density profiles of Cl^-^ (**A** & **D**), Na^+^ (**B** & **E**) and water (**C** & **F**), with ECC-rescaled ionic charges and a standard non-polarizable force field for all other atoms, in nanopores of different radii. ρ(r)/ρ_b_*A* represents the symmetrized number density, ρ(r), sampled from the hydrophobic region of the nanopore and normalized by bulk density, ρ_b_, and internal surface area, *A*, of the nanopore. The vertical dashed lines indicate the interface where the aqueous solution meets the wall of the nanopore (at radius R*_pore_*) colored accordingly for each respective nanopore size.

In the ECC simulations, Cl^-^ clearly accumulates at the hydrophobic nanopore wall and the extent of this phenomenon is approximately consistent across nanopore sizes, indicated by the relative height of the interfacial peaks (**Fig. 3A**). This suggests that in 0.5 M NaCl, the surface propensity of Cl^-^ has reached saturation at the interface and any additional ions entering the hydrophobic region contributed to the density of inner, more bulk-like regions. In contrast, no such interfacial ion effects are observed in the NP simulations (**Fig. 3D**). Instead, an increase in pore radius simply yields a gradual increase in Cl^-^ density, with the interfacial layer still devoid of ions.

Na^+^ is excluded from the hydrophobic interface and favors the middle of the pore in all nanopore sizes in both the ECC and NP simulations (**Fig. 3B & 3E**). Given that the smaller Na^+^ ions favor being fully solvated, the increase in pore radius, and thus the increase in volume of the bulk-like water regime, enables more Na^+^ to retain their hydration shells leading to an increase in number density. These effects are again comparable to those seen at aqueous/air and aqueous/decane interfaces (26, 35).

The water number density profiles remain similar for both NP and ECC simulations. In the smallest nanopore, the water profiles (**Fig. 3C & 3F**) suggest that water is packed more densely towards the axis of the nanopore, whereas in larger nanopores the density of water becomes progressively more bulk-like with the introduction of more annular rings of water as the pore radius increases. Similar water structure and packing inside CNTs has been observed in MD simulations and experimentally using Raman spectroscopy methods (72, 75).

Overall, these results suggest the preferential accumulation of chloride ions (alongside exclusion of sodium ions) close to the hydrophobic pore walls are effects due to surface electrostatic interactions involving the ion rather than as a consequence of confinement *per se*. The inclusion of polarization effects through ECC appears not only to affect the structure of water and ions inside the nanopore, but also to increase substantially the anion densities near the interfacial layer – an effect that is not observed with NP force fields.

### Energetics of ion permeation

The number density profiles indicate that in the central hydrophobic region of the model nanopore, Cl^-^ interact preferentially with the pore wall. To explore further the influence of these interactions on ion permeation, we estimated free energy profiles along the pore axis for the *R_pore_* = 0.70 nm nanopore (**Fig. 4**). We examined how the energetics of ion permeation were impacted by the different ion models. To this end umbrella sampling simulations were performed for both the ECC and NP forcefields to obtain symmetrized one-dimensional potentials of mean force (PMF) for both a Na^+^ and Cl^-^ Convergence analysis indicated the resulting PMF profiles for Cl^-^ had converged (i.e. taking ≲1 kJ/mol change between each fraction of time as a sign of convergence) for both ion parameter sets.

**Figure 4:**
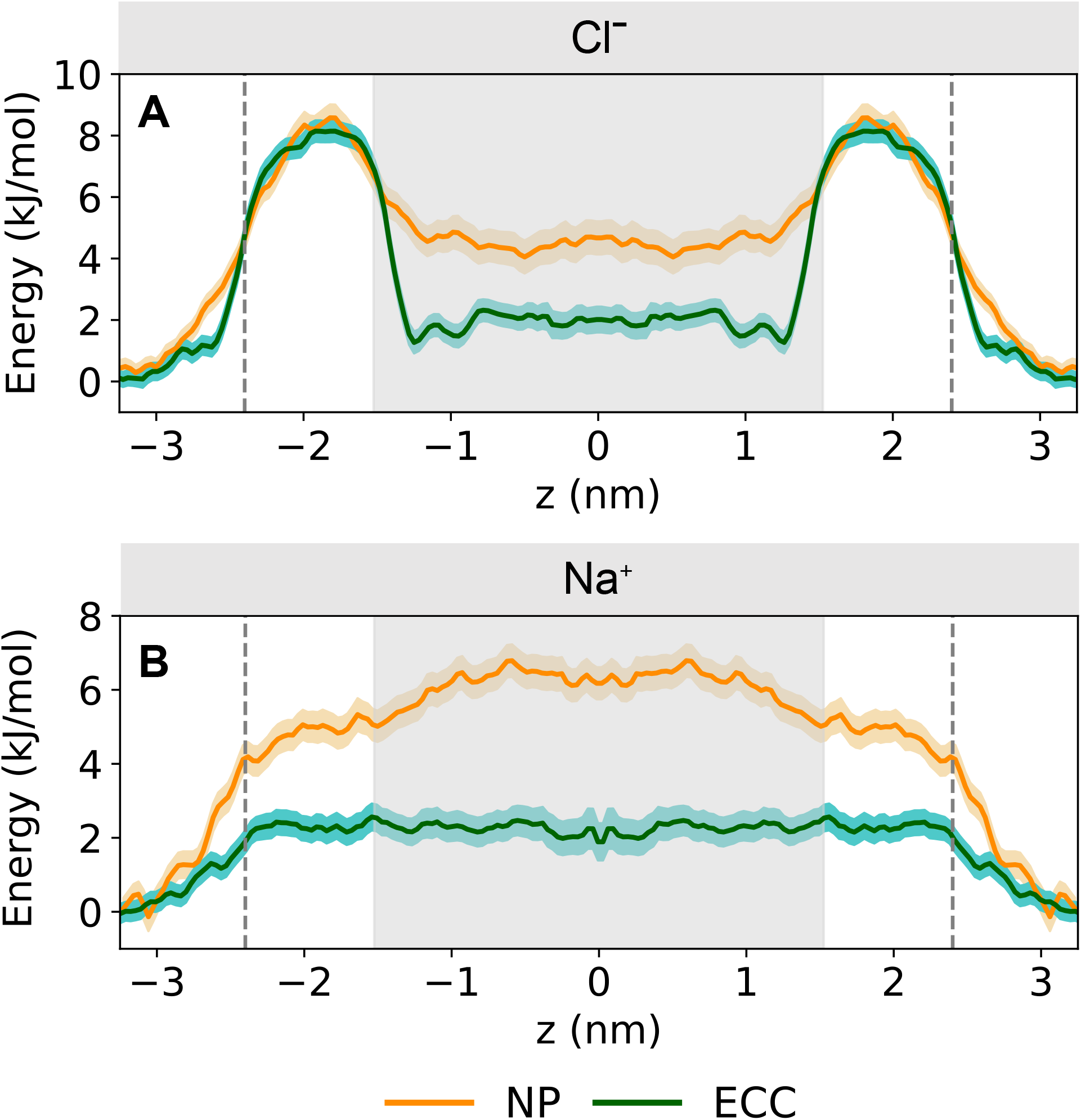
Single-ion PMFs profiles for **A** Cl^-^ and **B** Na^+^ permeating the model nanopore with ECC-rescaled ionic charges (green) and standard non-polarizable force field (yellow). The distance between the ion and the model nanopore center of mass is denoted by *z* where *z* = 0 represents the center of the pore. The solid lines indicate the free energy profile calculated from the final 5 ns of each umbrella window. Confidence bands were calculated by taking the standard error over independent 1 ns sampling blocks over the time period sampled. The dashed vertical lines denote the extent of the nanopore.

For Na^+^, both the NP and the ECC simulations yielded PMFs with a broad energetic barrier with a maximum in the center of the nanopore (i.e. at *z* = 0). This barrier height was ~ 6.2 kJ/mol for the NP PMF and was reduced ~ 3-fold relative to this (to ~ 2.0 kJ/mol) for the corresponding ECC PMF. Interestingly, simulations of a PMF for Na^+^ along the length of the channel formed by gramicidin A have shown a ~ 3-fold reduction in the central barrier height when comparing CHARMM27 with the AMOEBA polarizable forcefield (76) and a ~4-fold reduction comparing CHARMM27 with the CHARMM DRUDE polarizable forcefield (77).

The shape of the Cl^-^ PMFs is more complex, but overall is preserved between the NP and ECC simulations. In both cases there is an energy barrier of *~*8.3 kJ/mol at *z* = 2 nm corresponding to the narrow (radius ~ 0.4 nm; **Fig. 1D**) polar regions at the entrance to the pore. In this region, it is likely that the ions experience steric effects from the restrained water molecules. The energetic penalty in the polar region may additionally be due to the requirement for Cl^-^ to partially strip its solvation shell, releasing ~ 2 water molecules which are incompletely compensated for by less favorable interactions with the restrained water molecules which form the pore lining in this region (Fig. S2). The height of this barrier is comparable for the NP and ECC PMFs.

For both the NP and ECC Cl^-^ PMFs there is a broad energetic well centered around z = 0. However, for the NP simulations, this well is ~ +4.6 kJ/mol relative to solution outside the pore, whereas for the ECC simulations the difference (bulk to hydrophobic pore region) is ~ +1 kJ/mol at z = 1 nm rising to +2 kJ/mol in the center. Thus, at z = 1 nm (just inside the hydrophobic central region) Cl^-^ is stabilized nearly 4-fold in the ECC simulations relative to in the NP simulations. This correlates with previous studies (see discussion above) which have suggested that Cl^-^ is preferentially stabilized at a water/hydrophobic interface when ECC or a fully polarizable model are employed in simulations. It also agrees with the number density profiles above (**Fig. 2A & 3A**).

### Ion hydration within the nanopore

To examine the molecular origin of the differences in energetic profiles in more detail, we considered the changes in ion hydration at different radial locations inside the hydrophobic central region of the nanopore. The hydrophobic region was divided into four sections radially from the axis of the pore up to the pore wall in increments of 0.175 nm (**Fig. 5A**). The following analysis was performed on the final 5 ns of each simulation. For ions present in each region, radial distribution functions (RDFs) were computed between the ion and oxygens in the water molecules, i.e. *g_ION-O_*(*r*), for both the NP and ECC simulations. Ion-water coordination numbers were obtained by evaluating the cumulative ion-oxygen RDFs up to the first minimum, corresponding to the number of oxygens in the first hydration shell. All RDFs were calculated on systems solvated with 0.50 M NaCl solution and with nanopore radius of 0.70 nm.

**Figure 5:**
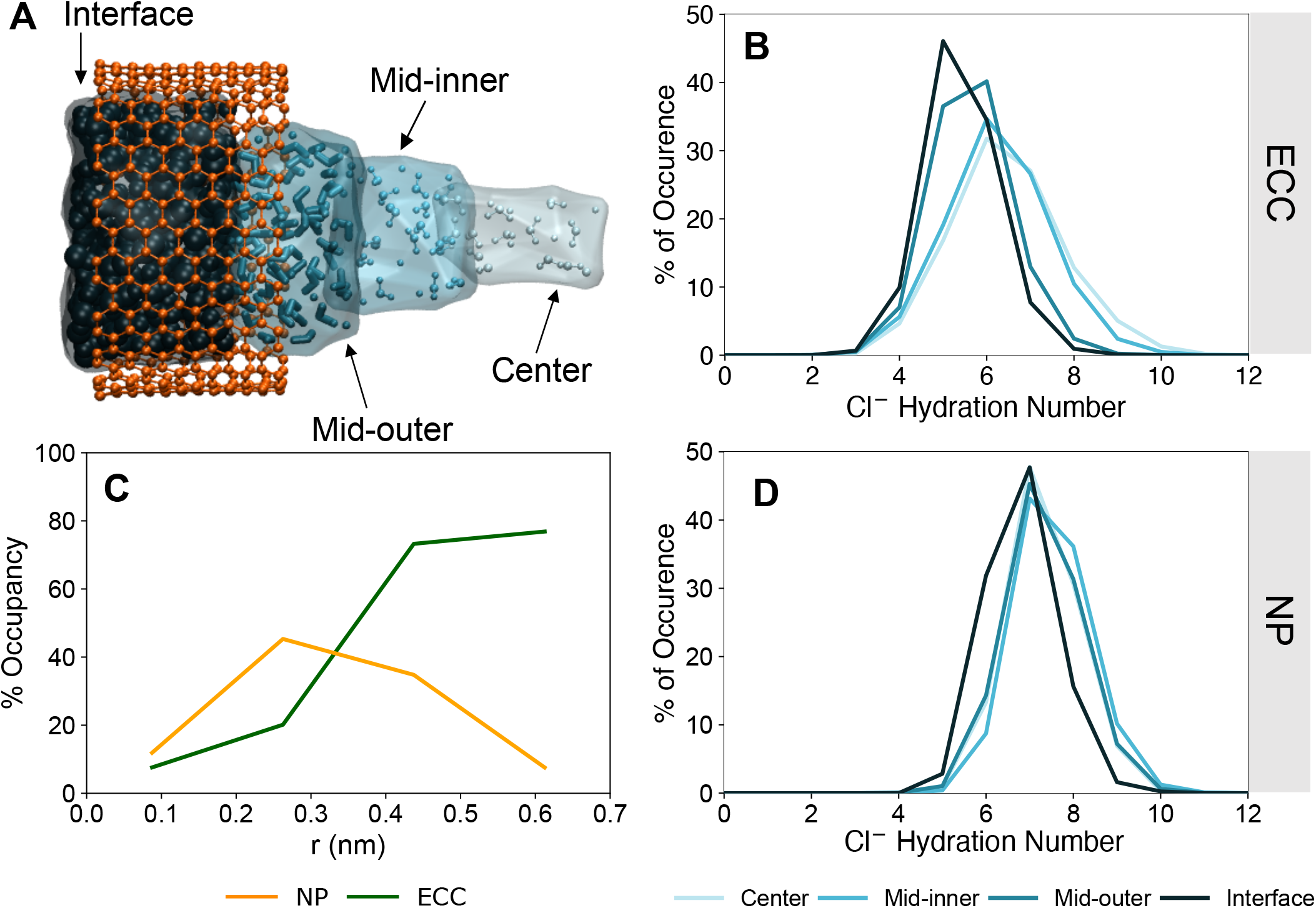
Cl^-^ hydration structure inside radial sections of the hydrophobic region of the pore. **A** Schematic of the hydrophobic region of the pore divided into four 0.175 nm radial sections colored in decreasing shades of blue. **B** & **D** The proportion of Cl^-^ with various hydration numbers in defined radial regions with the ECC method (**B**) and the non-polarizable force field (**D**). **C** shows the percentage occupancy of each radial section by Cl^-^. With ECC (green line), Cl^-^ spends a significantly greater proportion of time within the interfacial layer whereas with the NP force field (orange line) Cl^-^ tends to occupy regions away from the pore wall.

The Cl^-^ -O RDFs display a change in radius of the first hydration shell both between ion models and ion location. For ions outside the pore, the RDFs achieve the positions of first maxima for ECC and NP models at 0.33 nm (**Fig. S4B & S4C**) and 0.32 nm (**Fig. S4B & S4D**) respectively, whilst sharing relatively similar positions for the first minima. Density functional theory calculations report an RDF first peak distance between 0.31 and 0.32 nm which also agrees with experimental data (78, 79). Similar differences have been noted in simulations comparing Cl^-^ in bulk solution using NP and Drude polarizable force fields (80). RDFs with the ECC method show a reduction in peak intensity for the first peak which is associated with a difference in coordination number between force fields. Outside the nanopore (i.e. in bulk solution), the coordination number of chloride ions is 5.9 for the ECC compared to 7.1 for the NP force field. The ECC value falls within range of values 5.1 – 6.3 predicted by *ab initio* calculations (81) and 6.1 reported by first principles MD simulations (71). Earlier neutron scattering data report hydration numbers of 7.0 ± 0.4 (79) whilst other experiments yielded values of 5.5 ± 0.4 (82). Therefore, the coordination numbers for bulk electrolyte predicted in this study from the ECC simulations are consistent with previous theoretical and experimental data. (We note that our normalized RDFs for outside the pore do not reach a value of 1 at large distances.

This is due to the simulation box dimensions and the method of calculation which involves water-oxygen atoms from the whole system including near the bilayer-pore complex. Therefore, this is a non-homogeneous system).

The Cl^-^ first hydration number distributions indicated a significant shift towards lower values for ions at the interface of the pore (i.e., within 0.175 nm of the CNT pore wall), attaining an average value of ~ 5 with a significant fraction of ions with hydration numbers of ≤ 4.5 (**Fig. 5B**). In comparison, towards the (radial) center of the nanopore where water density is more bulk-like, the hydration number is on average ~ 6.1. Between the interface and radial center of the pore, the average hydration number ranges within these limits (**Fig. 5B**). The shape of the distributions become progressively broader with increasing distance away from the interface which indicates that the structure of the hydration shell becomes more flexible, approaching more bulk-like water behavior. In comparison, the Cl^-^ hydration shell is less flexible when using the NP force field. This is reflected in the hydration number distributions which remain tighter for NP (**Fig. 5D**) compared to ECC and are centered on an average coordination number of ~ 7.0 in all radial sections inside the nanopore, suggesting the hydration shell remains predominantly intact.

It is interesting also to consider the proportion of time Cl^-^ spends in various regions or, in other words, the percentage occupancy of each radial section (**Fig. 5C**). With ECC, Cl^-^ spend a significantly greater proportion of time in the interfacial layer whereas with the NP force field, Cl^-^ is more inclined to occupy regions away from the pore wall. These findings align with the number density distributions (**Fig. 2A & 3A**).

In contrast with Cl^-^, the RDFs for Na^+^ are in good agreement for inside compared to outside (i.e. bulk electrolyte) the pore for the first hydration shell with the ECC force field (**Fig. S3B**), suggesting that Na^+^ retains their solvation shell under hydrophobic confinement within these pores. The coordination number of Na^+^ at the interface is ~ 4.2, whereas outside the pore (i.e. in the bulk electrolyte) Na^+^ has a coordination number of ~ 4.8 when employing ECC (**Fig. S3A**). By contrast, a coordination number of ~ 5.0 is seen for Na^+^ regardless of location using the NP force field (**Fig. S3D**). Coordination numbers within the range 4.9-6.1 have previously been predicted for Na^+^ in bulk water using *ab initio* methods (81, 83). Older experiments of X-ray and neutron diffraction, and Raman spectroscopy predict coordination numbers between 4-8 for sodium in aqueous solution (84). The proportion of frames analyzed with Na^+^ present indicated that they were more likely to occupy the space away from the interface (**Fig. S3C & S3F**). These results reinforce the findings from number density calculations.

Overall, using the ECC parameters, Cl^-^ has a more flexible hydration shell and is more inclined to partially desolvate to favorably interact with the hydrophobic pore wall. Conversely, the Cl^-^ hydration shell remains predominantly intact using the NP force field and occupies regions away from the interface. Na^+^ prefers to remain more solvated by moving through the nanopore closer to the pore axis where the structure of water is more bulk-like relative to the pore wall.

### Robustness and sensitivity to the ECC model

The ECC model compensates for the absence of electronic polarization effects in condensed phase simulations using conventional fixed charge MD simulations. As discussed by Leontyev and Struchebrukhov (34, 85) this can be achieved by scaling of ionic charges. This approach has been investigated in a number of studies of ions at interfaces by Jungwirth *et al*. (20, 26, 86, 87). It has been shown to agree well with simulations of halide anions using a polarizable forcefield (88), and has been used in a number of simulation studies of lipid bilayer membranes and their interactions with ions (37, 89, 90).

There are some variations in the published ECC model (20, 26, 86, 87). Specifically, the more recent ECCR model (86) includes some additional small changes to ionic van der Waals parameters. We therefore compared ion density profiles as a function of radial position for our nanopore models using the ECC model in which only ionic charges have been modified and using the ECCR model in which van der Waals radii were also adjusted. As can be seen from **Fig. S5**, whilst the results show some sensitivity of the details of ion profiles to the model employed, in all cases for three different pore radii, the fundamental basic observation of accumulation of Cl^-^ ions alongside depletion of Na^+^ ions at the hydrophobic wall of the nanopore is observed. Furthermore, in all three pore models, for both the ECC and ECCR treatments, the difference in distribution of ions at the pore wall (calculated as Δ*r*(*Cl-Na*), i.e. the difference in distance from the nanopore wall at which the ionic concentration rises to 50%; see **Fig. S5** for details) is ~ 0.2 nm, i.e. the outermost solvation shell of the nanopore experiences an enhanced local concentration of anions. We are therefore confident that our observations are robust to reasonable variations in the implementation of the ECC model.

To explore this further, we have calculated free energy profiles using the ECCR model following the same simulation protocol as for the ECC and NP PMF calculations. As seen in **Fig. 4** (also see **Fig. S6**) the shape of the Cl^-^ PMFs are preserved between ECCR, ECC and NP simulations, exhibiting similar notable features corresponding to the polar regions at the pore entrances and a central barrier depletion in the hydrophobic core (z = 0 nm). For Cl^-^ in the polar regions, the energy barrier for ECCR is ~ 1 kJ/mol less compared to ECC and NP (**Fig. S6A**). This can be attributed to the additional van der Waals radius rescaling applied in ECCR and hence permeating ions likely experience fewer steric clashes. The free energy barrier for Cl^-^ using the ECCR parameters in the center of the nanopore (*z* = 0 nm) is ~ 4 kJ/mol which is a little lower than the NP energy at this location. This is perhaps somewhat unexpected given the number density profiles for Cl^-^ employing ECCR and ECC share similarities associated with the enhanced localization of anions to the hydrophobic pore interface (**Fig. S5**) as discussed previously. For Na^+^, an energetic maximum of ~ 3 kJ/mol is reached at *z* ~ 1 nm (**Fig. S6B**). where the ion begins to move into the hydrophobic core of the nanopore and away from the polar regions. The free energy at *z* = 0 nm is maintained at ~ 2.5 kJ/mol relative to outside the nanopore. Contrary to the pattern seen with Cl^-^, the free energy associated with ECCR for Na^+^ is closer to the PMF profile for ECC in the hydrophobic region which corresponds to the comparable number density plots in **Fig. S5**. However, for both ionic species the free energy of ions in the center of the pore follows the order NP > ECCR > ECC. Thus, this additional work highlights the sensitivity of nanoscale effects to ion parameters in MD simulations.

Additional exploration of ECC model sensitivity used an aqueous/decane interface as a simple model slab system (*cf*. (26)). We examined ECC sensitivity in this system to the water model employed, using four widely employed models (SPC/E, TIP3P, TIP4P and TIP4P/2005; see **Fig. S7A**). In all cases, local accumulation of Cl^-^ and depletion of Na^+^ at the hydrophobic interface was seen, but the details of the ion concentration vs. *r* profiles showed some sensitivity to the water model such that Δ*r*(*Cl-Na*) ranged from ~ 0.1 to ~ 0.2 nm. We also made preliminary comparisons with a polarisable model (AMOEBA (91, 92)) which revealed a comparable local accumulation of Cl^-^ /depletion of Na^+^ at the hydrophobic interface with Δ*r*(*Cl-Na*) = ~ 0.3 nm (**Fig. S7B**). This is in broad agreement with previous comparisons of ECC with polarisable models (20, 88).

Together these results suggest that our observation of local accumulation of Cl^-^ at the hydrophobic nanopore wall is robust to variations in the ECC model, and that this model is likely to mimic more computationally demanding polarizable simulations. It is also helpful to consider how well these simulations mimic experimental reality for anions close to a graphene-like hydrophobic surface. A recent study (93) compared anion adsorption to graphene/water interfaces as measured by surface-sensitive spectroscopy with (non-polarizable) MD simulations. This revealed that the experimental free energy of anion adsorption to a water/graphene interface could be reproduced by scaling the anion (iodide) charge by ~ 0.8, as is the case in the ECC model. This in turn suggests that the ECC model is likely to quantitatively reproduce local anion accumulation at the hydrophobic interface of a CNT-derived model nanopore.

## Conclusions

In this study, we have designed a model nanopore which mimics aspects of the critical pore region within pLGICs. This allows us to explore the interactions of ions and water in hydrophobically confined environments. Our results underline the importance of including polarization effects to model more accurately the interactions of ions (especially Cl^-^) with the hydrophobic surfaces that often line pores of this size. In particular, our findings demonstrate that when using the ECC model, the larger, more polarizable, Cl^-^ ions preferentially reside in the outermost interfacial layer of the hydrophobic region inside the nanopore, whereas the smaller, ‘ hard’ Na^+^ ions are repelled from the interface and occupy more bulk-like regions. Using this model, we investigated the effect of NaCl concentration and pore radius on ion localization and report that this trend persists. These observations resemble interfacial effects observed at aqueous/air and aqueous/oil (**Fig. 6B**) interfaces and are not reproduced using the non-polarizable force field (26, 35).

**Figure 6:**
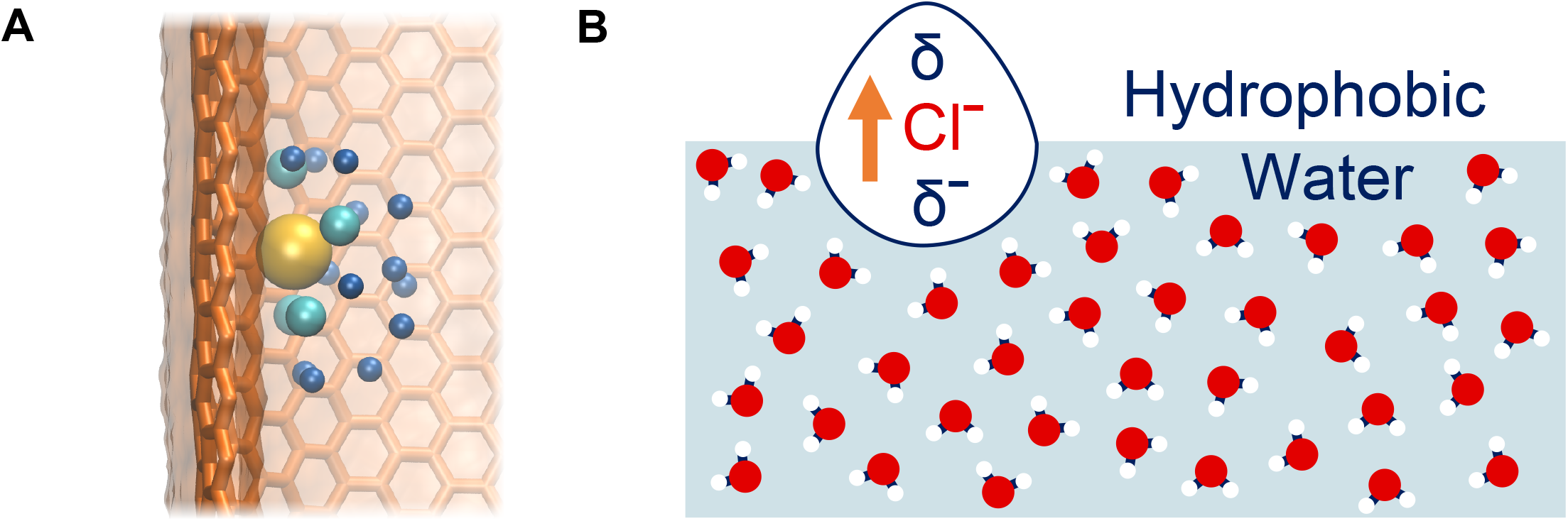
**A** Snapshot of Cl^-^ partially desolvating to favorably interact with the internal hydrophobic interface of the model nanopore. Cl^-^ is represented in yellow and the oxygen atoms of the water molecules from the first and second hydration shells are represented by light and dark blue beads respectively. **B** Schematic diagram of an induced dipole in Cl^-^ at a hydrophobic/water interface. Adapted from (25).

Through analysis of the ion hydration structure inside the hydrophobic core, it has been possible to investigate ion solvation as a function of radial position from the pore axis. A Cl^-^ in proximity of the hydrophobic pore wall can be seen to partially lose its hydration shell to form favorable interactions with the hydrophobic wall of the pore (**Fig. 6A**). This behavior is not captured without inclusion of polarization effects. Conversely, as Na^+^ ions dominantly occupy bulk-like regions, their solvation shell remains largely intact.

Free energy profiles for ions correlate with their number density profiles. The PMF profile for Cl^-^ with ECC suggests the partial loss of its hydration shell is energetically advantageous whereas the barrier for sodium is higher. Therefore, this suggests that the hydrophobic core of the nanopore exhibits a degree of selectivity for Cl^-^ over Na^+^. Comparable Cl^-^ hydrophobic interactions have recently been reported in biological Cl^-^ channels and transport proteins, for example in the NTQ Cl^-^ - pumping rhodopsin (PDB: 5G28) (94) and in the bestrophin-1 chloride channel (PDB: 4RDQ) (95). Similarly, some synthetic anionphores (biotin[6]uril hexaesters) exploit C—H hydrogen bond donors to selectively transport softer anions over harder, more basic anions (96). Thus, it is important to model accurately the interactions of anions with hydrophobic binding sites in channels, transporters, and synthetic carriers. In this context, it is of interest that Orabi *et al*. (33) have explored the effect of modifying ion van der Waals parameters using NBFIX to mimic polarizability effects through pair-specific LJ parameters to override Lorentz-Berthelot combination rules in simulations of a CLC Cl^-^ transporter. Their results demonstrated that with the standard CHARMM36m force field, Cl^-^ experienced dissociation from the binding site observed in the crystal structure of the protein, whereas the ion remains bound with the NBFIX parameters. Taken together, these studies indicate that further investigation is required into how the inclusion of electronic polarizability in simulations may influence our understanding of anion behavior in both synthetic and biological anion selective structures.

Overall, our analysis of a model biomimetic nanopore reveals contrasting ion behavior that may provide insights into the fundamental principles of anion selectivity and has the potential to influence technological applications. Our findings also suggests that the inclusion of electronic polarizability in ion modelling is key to accurately capturing Cl^-^ behavior. Moreover, this current study contributes to the longstanding debate over force field accuracy and whether more explicit treatment of electrostatics is necessary at the expense of computational efficiency (97, 98). More extensive simulations comparing interactions between Cl^-^ and hydrophobic interfaces with the use of explicitly polarizable force fields may provide new mechanistic insights into high anion selectivity.

## Acknowledgements

This work was supported by grants from the EPSRC and BBSRC and by an EPSRC iCASE studentship award (to LXP) in collaboration with IBM Research. It was also supported by the Hartree National Centre for Digital Innovation - a collaboration between the Science and Technologies Facilities Council and IBM.

## Notes

### Competing Interest Statement

The authors have declared no competing interest.

