## Supplementary Information for "Influence of effective polarization on ion and water interactions within a biomimetic nanopore"

*Supplemental Information*

Figure S1

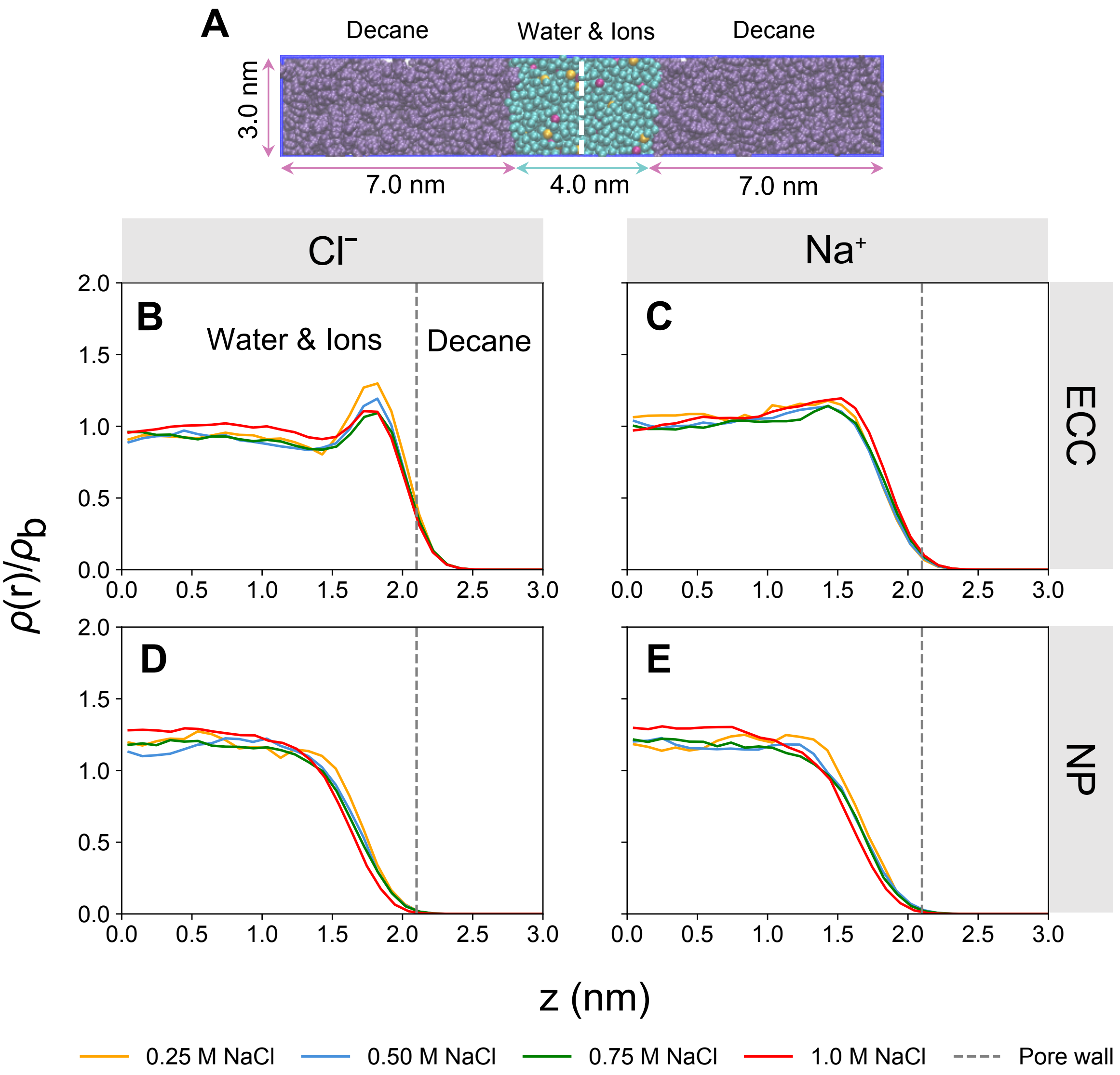


***Figure S1*****:** **A** Schematic of the aqueous/decane interface as a model slab system. The slab system consists of decane (purple), water (pale blue), $Na^{+}$ (pink) and $Cl^{-}$ (yellow). Symmetrized number density profiles are shown for $Cl^{-}$ (**B** & **D**) and $Na^{+}$ (**C** & **E**) at the aqueous/decane interface with ECC-rescaled charges and a non-polarizable force field at various salt concentrations. $\rho(r)/\rho_b$ represents the symmetrized number density, $\rho(r)$, which has been normalised by bulk density, $\rho_b$. The variable *z* is the distance from the center of the simulation box and the vertical dashed line corresponds to the aqueous/decane interface.

Figure S2

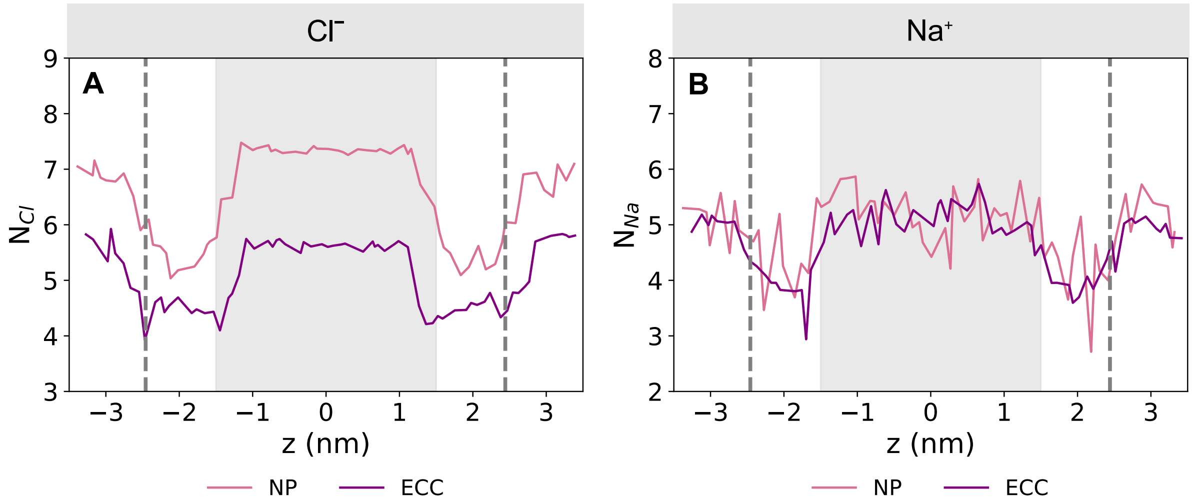


***Figure S2:*** Variation in the first hydration shell of $Cl^-$ (**A**) and $Na^+$ (**B**) as a function of *z*, the distance between the ion and the model nanopore center of mass. $N_{Cl}$ and $N_{Na}$ denote the coordination number of water oxygen atoms in the first hydration shell of $Cl^-$ and $Na^+$ respectively, with ECC-rescaled ions (purple) and a non-polarizable (NP) force field (pink). The dashed vertical lines denote the extent of the nanopore.

Figure S3

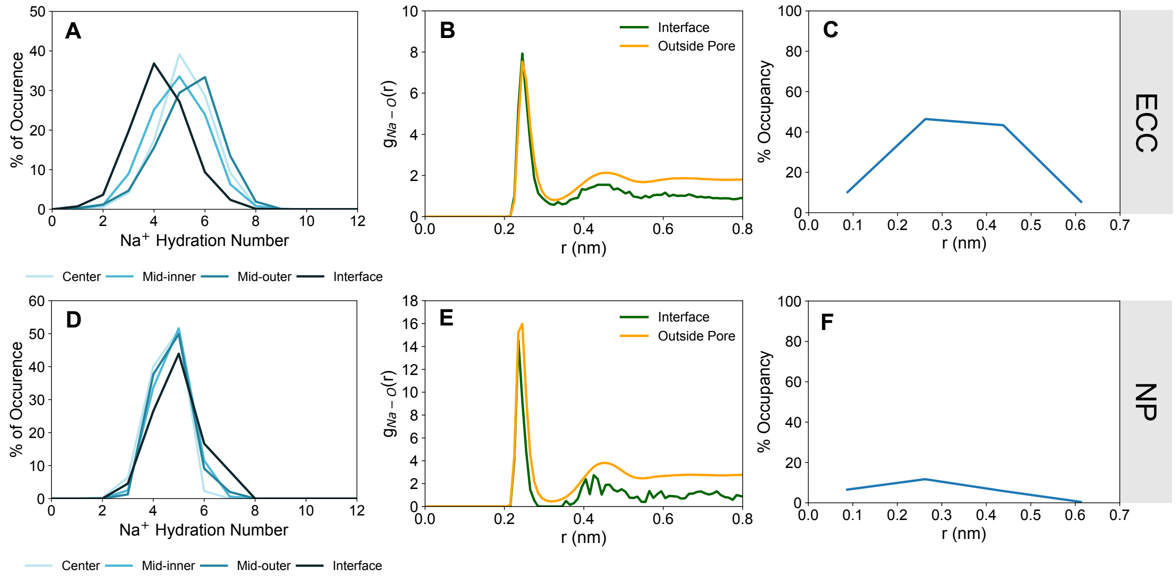


***Figure S3:*** $Na^+$ hydration structure inside radial sections of the hydrophobic region of the pore with the ECC method and a non-polarizable (NP) force field. **A** & **D** show the proportion of $Na^+$ with various hydration numbers in defined radial regions for ECC and NP respectively. **B** & **E** show the radial distribution function of water oxygen atoms around $Na^+$ in the interfacial layer inside the pore (green) and outside the pore in bulk solution (yellow) for ECC and NP respectively. **C** & **F** show the percentage occupancy of each radial section by $Na^+$. With both ECC and the NP force field, $Na^+$ tend to occupy regions away from the hydrophobic pore interface.

Figure S4

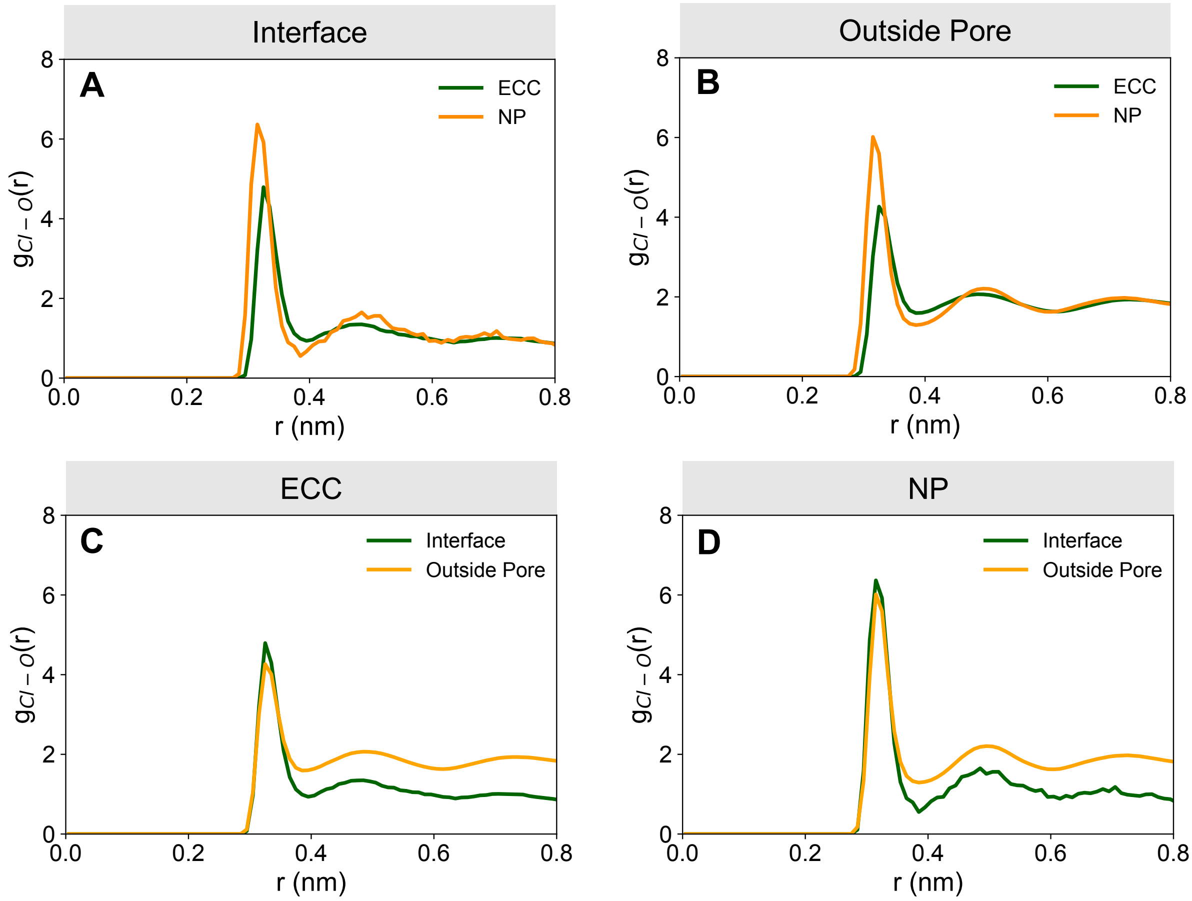


***Figure S4:*** Radial distribution function (RDF), $g_{Cl-O}$(r), of water oxygen atoms around $Cl^{-}$ at the interface (**A**) and outside the pore (**B**) with ECC and standard non-polarizable (NP) force field. **C** & **D** show the same RDFs as a comparison between the interface and outside the pore for a given force field parameter set.

Figure S5

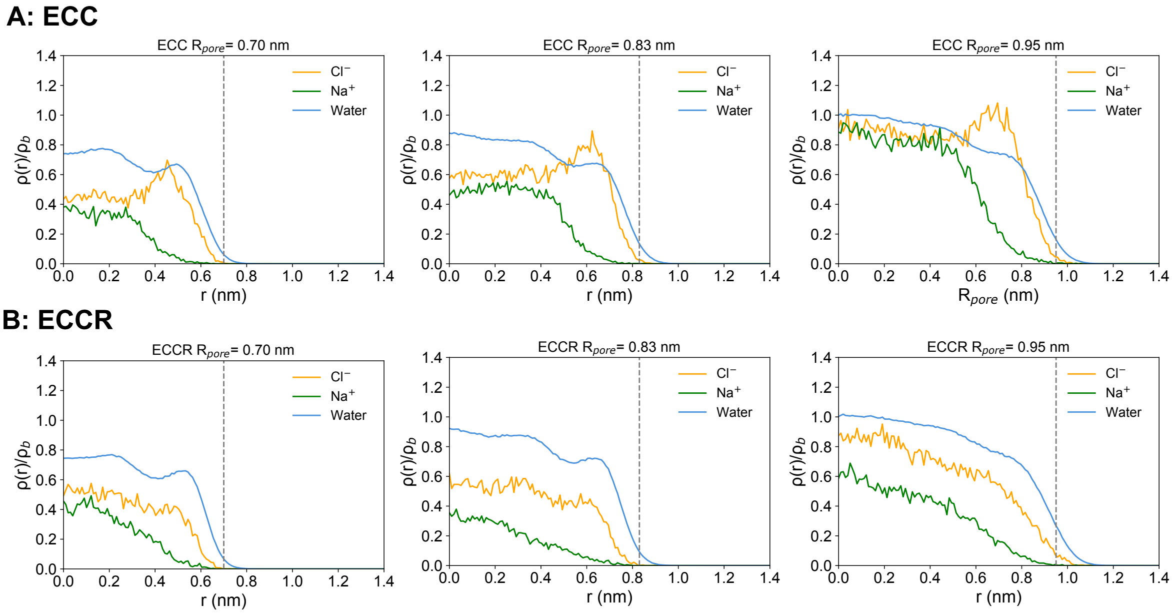


***Figure S5:*** Symmetrized number density ($\rho(r)/\rho_b$) profiles of $Cl^-$, $Na^+$ and water, with ECC-rescaled ionic charges (**A**) and ECCR (**B**) (ECC with an additional small van der Waals radii rescaling) in nanopores of different radii. The variable r is the radius of the nanopore which extends from 0 (pore axis) to $R_{pore}$ (the interface where the salt solution meets the wall of the nanopore). The grey vertical dashed lines represent $R_{pore}$.

Figure S6

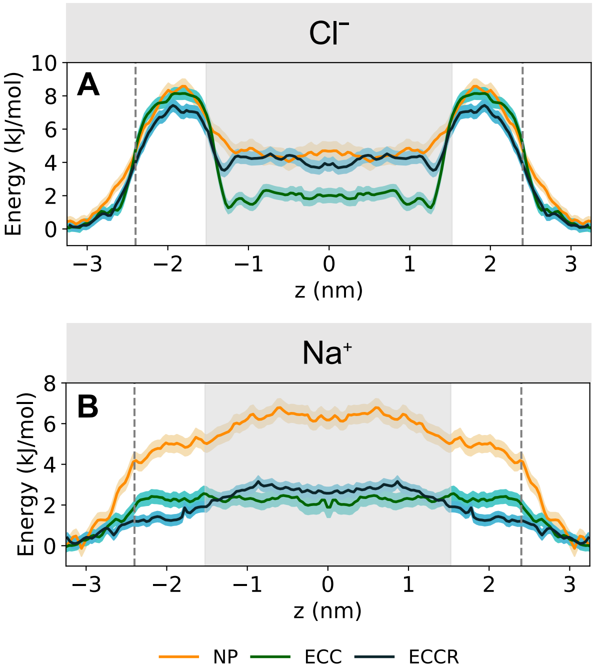


***Figure S6:*** Single-ion PMF profiles for **A** $Cl^-$ and **B** $Na^+$ permeating the model nanopore with ECC-rescaled ionic charges (green), standard non-polarizable (NP) force field (yellow) and ECCR (ECC with an additional small van der Waals radii rescaling) (dark blue). The distance between the ion and the model nanopore center of mass is denoted by $z$ where $z = 0$ represents the center of the pore. The solid lines indicate the free energy profile calculated from the final 5 ns of each umbrella window. Confidence bands were calculated by taking the standard error over independent 1 ns sampling blocks over the time period sampled. The dashed vertical lines denote the extent of the nanopore.

Figure S7

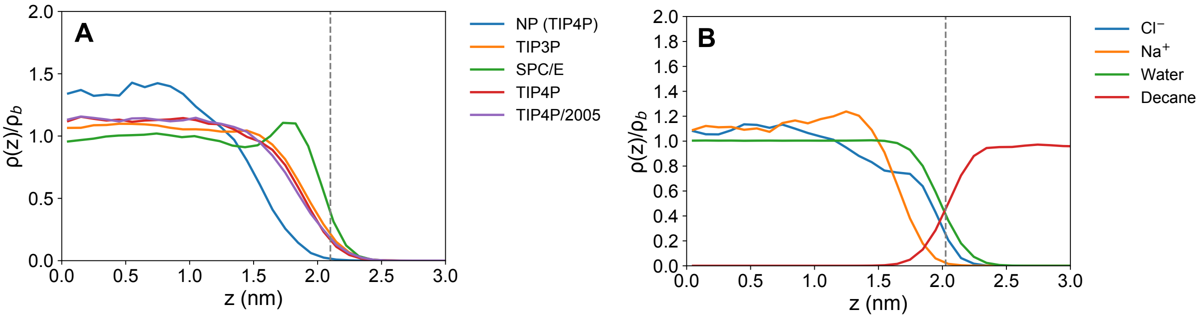


***Figure S7:*** Symmetrized number density ($\rho(r)/\rho_b$) profiles for $Cl^-$ (**A**) at an aqueous/decane interface employing various water models with ECC-rescaled charges and a non-polarizable (NP) force field using TIP4P water model. **B** shows the symmetrized number density profiles for $Cl^-$, $Na^+$ and water using the AMOEBA polarizable force field with the AMOEBA03 water model. The variable $z$ is the distance from the center of the simulation box and the vertical dashed line denotes the aqueous/decane interface.

Figure S8

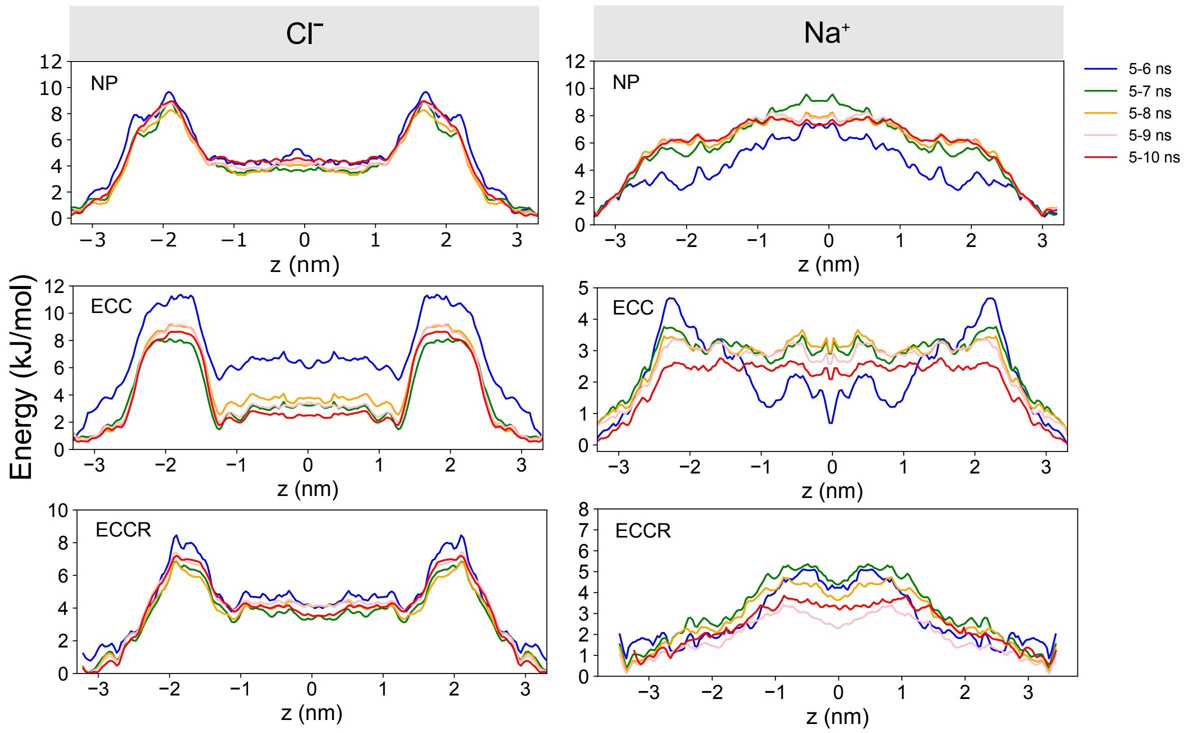


***Figure S8:*** Single-ion PMFs convergence analysis for $Cl^-$ and $Na^+$ using a non-polarizable (NP) force field, ECC and ECCR ion parameters. Convergence analysis was performed by calculating 1 ns cumulative sampling blocks over the sampling time (last 5 ns of simulation).
